# Identification of ER/SR resident proteins as biomarkers for ER/SR calcium depletion in skeletal muscle cells

**DOI:** 10.1101/2021.12.29.474463

**Authors:** Lacey K. Greer, Katherine G. Meilleur, Brandon K. Harvey, Emily S. Wires

## Abstract

Aberrations to endoplasmic/sarcoplasmic reticulum (ER/SR) calcium concentration can result in the departure of endogenous proteins in a phenomenon termed exodosis. Redistribution of the ER/SR proteome can have deleterious effects to cell function and cell viability, often contributing to disease pathogenesis. Many proteins prone to exodosis reside in the ER/SR via an ER retention/retrieval sequence (ERS) and are involved in protein folding, protein modification, and protein trafficking. While the consequences of their extracellular presence have yet to be fully delineated, the proteins that have undergone exodosis may be useful for biomarker development. Skeletal muscle cells rely upon tightly coordinated ER/SR calcium release for muscle contractions, and perturbations to calcium homeostasis can result in myopathies. Ryanodine receptor type-1 (RYR1) is a calcium release channel located in the SR. Mutations to the *RYR1* gene can compromise calcium homeostasis leading to a vast range of clinical phenotypes that include hypotonia, myalgia, respiratory insufficiency, ophthalmoplegia, fatigue and malignant hyperthermia (MH). There are currently no FDA approved treatments for RYR1-related myopathies (RYR1-RM). Here we examine the exodosis profile of skeletal muscle cells following ER/SR calcium depletion. Proteomic analysis identified 4,465 extracellular proteins following ER/SR calcium depletion with 1,280 proteins significantly different than vehicle. A total of 54 ERS proteins were identified and 33 ERS proteins significantly increased following ER/SR calcium depletion. Specifically, ERS protein, mesencephalic astrocyte-derived neurotrophic factor (MANF), was elevated following calcium depletion, making it a potential biomarker candidate for human samples. Despite no significant elevation of MANF in plasma levels among healthy volunteers and RYR1-RM individuals, MANF plasma levels positively correlated with age in RYR1-RM individuals, presenting a potential biomarker of disease progression. Selenoprotein N (SEPN1) was also detected only in extracellular samples following ER/SR calcium depletion. This protein is integral to calcium handling and *SEPN1* variants have a causal role in SEPN1-related myopathies (SEPN1-RM). Extracellular presence of ER/SR membrane proteins may provide new insight into proteomic alterations extending beyond ERS proteins. Pre-treatment of skeletal muscle cells with bromocriptine, an FDA approved drug recently found to have anti-exodosis effects, curbed exodosis of ER/SR resident proteins. Changes to the extracellular content caused by intracellular calcium dysregulation presents an opportunity for biomarker development and drug discovery.

## Introduction

The endoplasmic/sarcoplasmic reticulum (ER/SR) is the main reservoir for intracellular calcium [1], serving an imperative role for muscle contraction and relaxation. This extensive tubular network is also responsible for a myriad of cellular functions including, protein synthesis [2], protein modification [2], protein degradation [3], lipid metabolism [4], carbohydrate metabolism [5], and xenobiotic detoxification [6]. The SR is a specialized extension of the ER and coordinates the release of calcium during muscle contraction [7]. Depletion of ER/SR calcium causes the secretion of ER/SR resident protein in phenomenon described as exodosis [8]. Under physiological conditions, resident proteins are retained within ER/SR via a carboxy-terminal ER retention/retrieval sequence (ERS) that interacts with the KDEL receptor retrieval pathway [9]. Retention of ERS proteins is vital to cellular function, as many ERS proteins contribute to proper protein folding, modification, and trafficking [9]. Using a bioluminescent reporter protein, exodosis was recently shown to occur in skeletal muscle cells in response to pharmacologically induced ER/SR calcium depletion [10]. The presence of ERS proteins in the extracellular environment may be useful for identifying disruption to ER/SR calcium homeostasis and proteostasis in myopathies.

Ryanodine receptor isoform-1 (RYR1) is the predominant skeletal muscle calcium release channel located on the terminal cisternae of the SR and important for excitation-contraction coupling [11, 12]. Mutations in the *RYR1* gene have been implicated in a variety of rare congenital myopathies that include, multi-minicore disease (MmD), central core disease (CCD), and to a lesser extent, centronuclear myopathy (CNM), congenital fiber-type disproportion (CFTD), and King Denborough syndrome [11]. As of 2011, the prevalence of congenital myopathies due to *RYR1* mutations is reported to be 1:90,000 in the United States [13]. Moreover, *RYR1* variants can cause malignant hyperthermia (MH) susceptibility, a hypermetabolic response characterized by chronic muscle contraction and elevated body temperature in response to halogenated anesthetics [14, 15]. *RYR1* mutations have been implicated in chronic SR calcium leak, decreased RYR1 protein levels, and altered sensitivity [16–19]. There is currently no FDA-approved treatment for RYR1-related myopathies (RYR1-RM), although it is worth noting the RYR1 antagonist, dantrolene, is approved for use during MH events to attenuate excessive ER/SR calcium release [20].

There is an incentivized need to expedite orphan drug development and drug repurposing for the therapeutic intervention of rare diseases [21]. Henderson and colleagues recently screened 9,501 compounds approved by U.S. and numerous international regulatory agencies and characterized several for their anti-exodosis properties [10]. Given the propensity for SR calcium leak in *RYR1* variants, we sought to examine skeletal muscle cells for exodosis following ER/SR calcium depletion. Herein we describe a proteomic approach to identify potential biomarkers reflective of calcium dysregulation and investigate anti-exodosis effects of bromocriptine, an FDA-approved drug recently shown to have anti-exodosis properties.

## Results

### Calcium dysregulation elicits exodosis in human skeletal muscle cells

Previously, disruption of ER/SR calcium in primary skeletal muscle elicited the secretion of GLuc-SERCaMP, an exogenous reporter of exodosis. Curious if this observation extended to endogenous ERS proteins, human skeletal muscle cells were treated with 100 nM of thapsigargin (Tg) or vehicle for 8 hours. Tg is a pharmacological inhibitor of SR calcium ATPase (SERCA) commonly used to deplete ER/SR calcium reservoir. Tg increased general extracellular protein content (Supplemental Figure 1A) as well as a subset of ERS proteins previously shown to undergo exodosis [8], including mesencephalic astrocyte derived neurotrophic factor (MANF), protein disulfide isomerase (PDI), and liver carboxylesterases 1 and 2 (CES1 and CES2) (Figure 1A-C).

**Figure 1.**
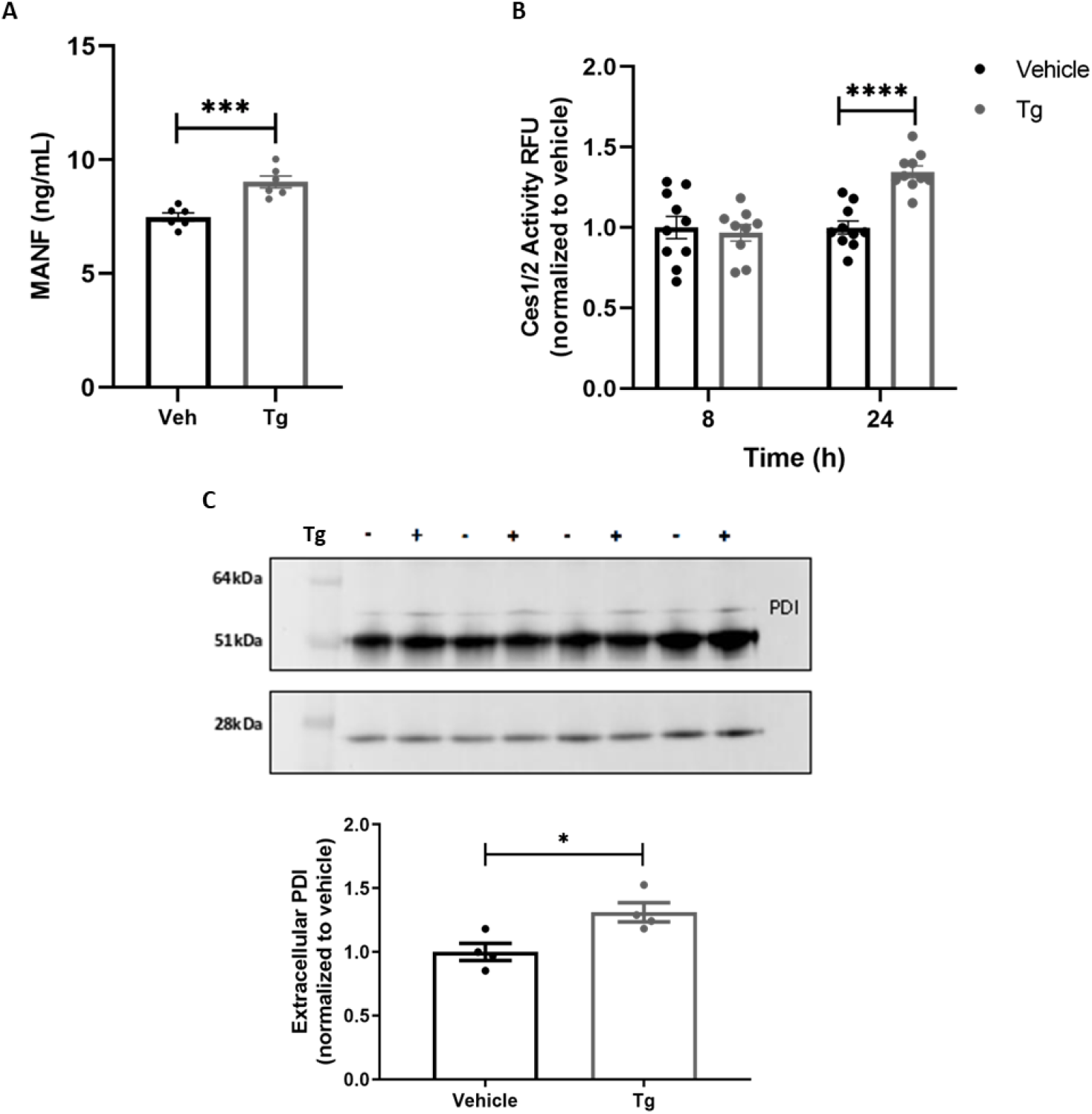
Targeted validation of exodosis-related proteins. A) MANF HTRF assay of media collected from T0034 skeletal muscle cell line pre-treated with dantrolene or bromocriptine 30 minutes prior to 100nM Tg for 8 hours. Extracellular MANF is increased following Tg treatment. Dantrolene or bromocriptine pre-treatment attenuate MANF exodosis, mean ± SEM, n= 12 wells/treatment group, ***p= 0.0008, unpaired two-tailed t-test. B) Carboxylesterase 1/2 fluorescence activity measured in media collected from T0034 skeletal muscle cell line treated with 100nM Tg for 8 and 24 hours, mean ± SEM, n=10/treatment group, ****p<0.0001, 2-way ANOVA, Sidak’s multiple comparison test. C) P4HB IP of media collected from T0034 skeletal muscle cell line treated with 100nM Tg for 8 hours, mean ± SEM, n=4/treatment group, *p=0.022, unpaired two-tailed t-test.

Next, we sought to use a more quantitative proteomics approach to identify additional extracellular ERS proteins by mass spectrometry (Figure 2). Of the 4,465 extracellular proteins identified, 1,280 proteins significantly changed following Tg treatment. 54 corresponded to ERS proteins [8, 9], with 33 ERS proteins significantly increased after Tg (Figure 2; Supplemental Figure 1B). Proteins identified by mass spectrometry corroborated findings in Figure 1. Functional analysis of the extracellular proteins identified by mass spectrometry are associated with protein binding (Figure 2, inset). Protein binding includes molecular chaperone function, gene transcription, protein translation, drug pharmacokinetics, and drug pharmacodynamics, suggesting that ER/SR calcium dysregulation poses numerous cellular challenges to muscle function. These data show ER/SR calcium dysregulation alters skeletal muscle proteome, eliciting the secretion of crucial ER/SR resident proteins whose intracellular absence and extracellular presence may impair cellular function.

**Figure 2.**
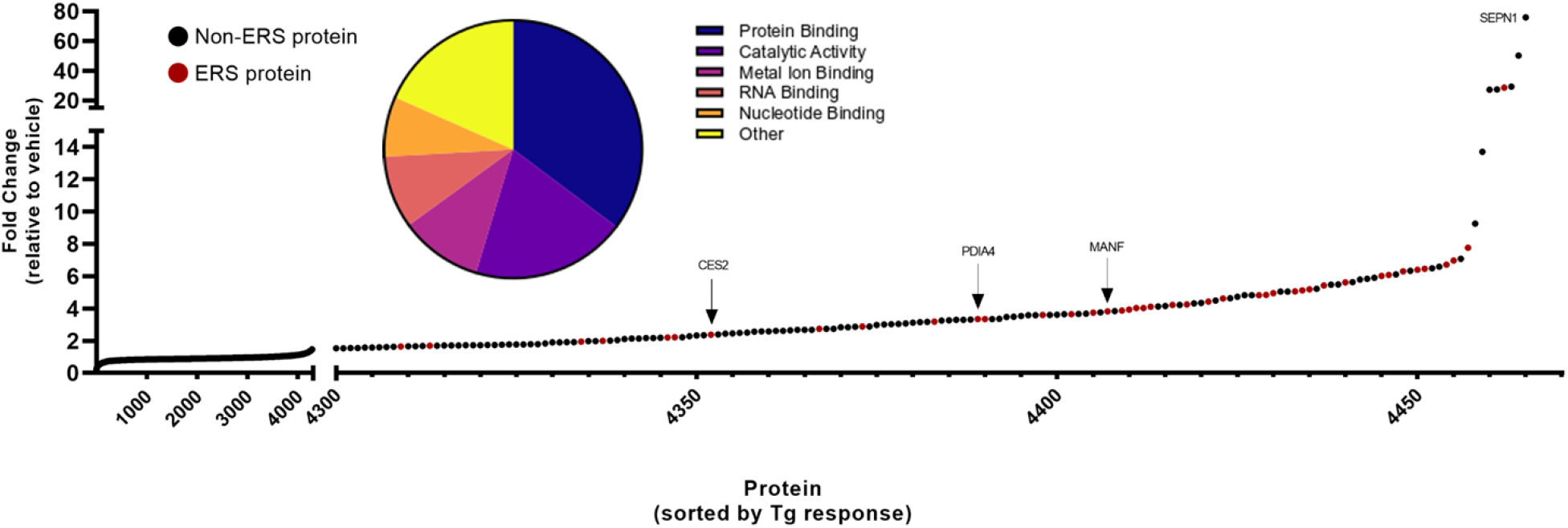
Calcium depletion elicits exodosis in skeletal muscle cell line. A) Mass spectrometry analysis of concentrated media from T0034 skeletal muscle cell line treated with 100nM Tg for 8 hours identified 4,465 extracellular proteins. Red circles indicate the 54 ERS proteins that were identified. Biological processes of all identified extracellular proteins highlighted by pie chart inset.

In addition to ERS proteins, several non-ERS proteins were also released from the cell in response to ER/SR calcium depletion. Of note, selenoprotein N (SEPN1) was the most abundant non-ERS extracellular protein detected based on previously established ERS criteria [8, 9] (Figure 2). Another example from our mass spectrometry data of a non-ERS protein with increased extracellular content is nucleotide exchange factor SIL1. SIL1 is a co-factor of ERS protein, HSPA5 (BiP) which displayed a 5.6115-fold increase following ER/SR calcium depletion (Supplemental Figure 1B).

### Bromocriptine attenuates exodosis in skeletal muscle cells

Variants within the *RYR1* gene have a causative role in the development of malignant hyperthermia (MH), a rare, but serious condition characterized by sustained muscle contraction, hyperthermia, and rhabdomyolysis in response to halogenated anesthetics [15]. Dantrolene is a RYR antagonist that stabilizes receptor resting state by increasing Mg^2+^ affinity and is currently the only FDA approved drug for MH, although a RYR1-selective inhibitor, 6,7-(methylenedioxy)-1-octyl-4-quinolone-3-carboxylic acid, was recently shown to be effective in murine MH models [20, 22]. Henderson et al, 2021 recently described a high throughput screen using the exodosis reporter *Gaussia* luciferase-secreted ER calcium modulated protein (GLuc-SERCaMP) and identified several FDA approved drugs effective in attenuating exodosis including dantrolene [10]. Of the additional compounds identified, bromocriptine, verapamil, dextromethorphan, and diltiazem partially attenuated SERCaMP release in primary skeletal muscle cells pre-treated for 30 minutes prior to 8-hour 100nM Tg treatment [10]. Moreover, bromocriptine consistently displayed anti-exodosis effects in models of stroke and Wolfram syndrome, prompting us to test its ability to reduce extracellular ERS proteins in skeletal muscle cells [10]. Towards this, we pre-treated skeletal muscle cells with 20μM bromocriptine or 50μM dantrolene 30 minutes prior to Tg treatment. No difference in overall extracellular protein content was observed among pre-treatment with 20μM bromocriptine or 50μM dantrolene (Figure 3A). However, pre-treatment with 20μM bromocriptine for 30 minutes prior to Tg-induced ER/SR calcium depletion significantly reduced the amount of extracellular ERS proteins (Figure 3B). Dantrolene pre-treatment exhibited a modest attenuation of ERS proteins, albeit to a lesser extent than bromocriptine (Figure 3B; Supplemental Figure 2A, B). One ERS protein, MANF, was increased in media evidenced by mass spectrometry data, western blot, and homogenous time-resolved fluorescence (HTRF) assays (Figure 1A; Figure 2; Figure 3D, E; Supplemental Figure 1B, 2B). Increased secretion of MANF has been previously observed during exodosis [8, 10]. Extracellular MANF increased approximately 3.8-fold upon ER/SR calcium depletion in skeletal muscle cell line (Supplemental Figure 1B). Pre-treatment with 20μM bromocriptine for 30 minutes prior to 100nM Tg for 8 hours significantly reduced extracellular MANF levels (Figure 3C-E). Surprisingly, dantrolene did not attenuate extracellular MANF levels despite previously reported attenuation in SH-SY5Y neuroblastoma cell line, suggesting varying cell-specific responses (Figure 3C-E) [8].

**Figure 3.**
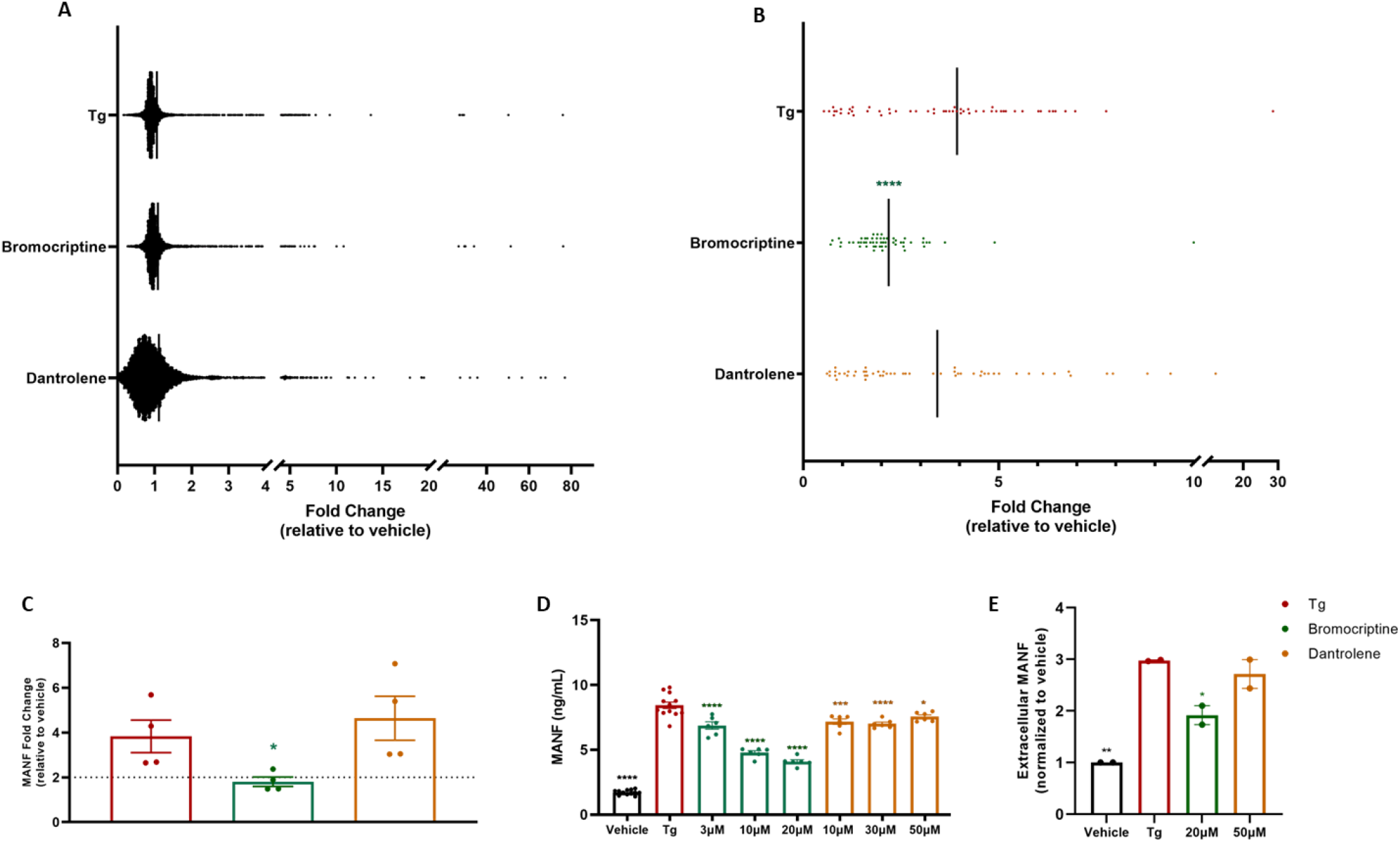
Bromocriptine attenuates exodosis in skeletal muscle cell line. A) Mass spectrometry analysis of media from T0034 skeletal muscle cell line pre-treated with vehicle, 20μM bromocriptine, or 50μM dantrolene 30 minutes prior to 100nM Tg for 8 hours. Solid line represents mean fold change B) Extracellular ERS proteins identified from mass spectrometry analysis pre-treated with vehicle, 20μM bromocriptine, or 50μM dantrolene 30 minutes prior to 100nM Tg for 8 hours; solid line represents mean fold change, ****p<0.0001, 1-way ANOVA, Dunnett’s multiple comparison test, Tg vs bromocriptine or dantrolene. C) Extracellular MANF identified by mass spectrometry in media from skeletal muscle cells pre-treated with vehicle, 20μM bromocriptine, or 50μM dantrolene 30 minutes prior to 100nM Tg for 8 hours; mean ± SEM, *p<0.05, 1-way ANOVA, Dunnett’s multiple comparison test. D) Extracellular MANF identified by MANF HTRF assay of media collected from T0034 skeletal muscle cell line pre-treated with vehicle, 3μM, 10μM, 20μM bromocriptine, or 10μM, 30μM, 50μM dantrolene 30 minutes prior to 100nM Tg for 8 hours; mean ± SEM, n= 6-12 wells/treatment group, *p<0.05, ***p<0.001, ****p<0.0001, 1-way ANOVA, Dunnett’s multiple comparison test, Tg vs other treatment groups. E) Densitometry analysis of western blot of concentrated media collected from T0034 skeletal muscle cell line pre-treated with vehicle, 20μM bromocriptine, or 50μM dantrolene, mean ± SEM, n=2/treatment groups, *p<0.05, **p<0.01, 1-way ANOVA, Dunnett’s multiple comparison test vehicle/Tg vs vehicle/vehicle, bromocriptine/Tg, or dantrolene/Tg.

### MANF is elevated in aging RYR1-RM-affected individuals

Given that Tg-induced ER/SR calcium depletion of human skeletal cell line elicited elevated extracellular MANF levels, we next sought to determine if circulating MANF levels were upregulated in RYR1-RM-affected individuals. Homogenous time-resolved fluorescence (HTRF) technology permits the measurement of biomarkers and analytes in the nanogram range from cell culture as well as human CSF, sera, blood, and plasma [23]. Commercially available MANF HTRF was used to assess levels of circulating MANF from human plasma samples obtained from a clinical trial NCT02362425 prior to intervention [24]. While no significant difference was observed among healthy volunteers and RYR1-RM-affected individuals, there was a significant correlation between circulating MANF levels and increased age observed only in RYR1-RM-affected individuals (Figure 4A-C). No correlation was detected between genotype or sex and MANF levels (data not shown). Collectively, our data support MANF as a biomarker of exodosis, with potential use for RYR1-RM progression and therapeutic intervention. Additional ERS proteins were identified and may serve as biomarkers of disrupted ER/SR calcium homeostasis and altered proteostasis in skeletal muscle cells, although further investigation is warranted.

**Figure 4.**
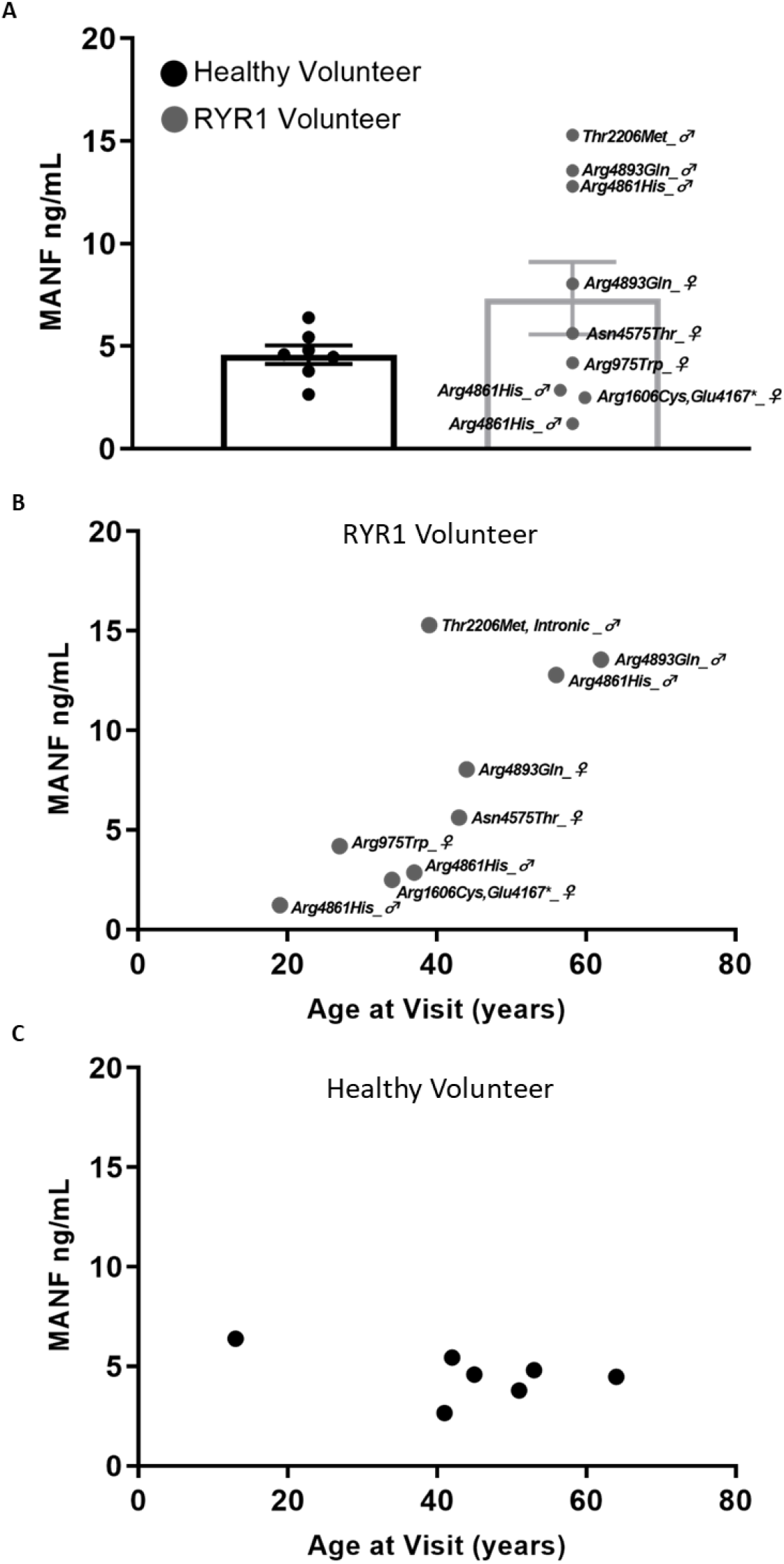
MANF is marginally elevated in individuals with *RYR1* mutations. A) Extracellular human MANF in plasma obtained from healthy volunteers or individuals with *RYR1* mutations, mean ± SEM, n=7 healthy volunteers, n=9 *RYR1* mutations, participant sex indicated next to confirmed genotype, p=0.1665, two-tailed t-test. B, C) Circulating levels of MANF is significantly correlated with age in individuals with *RYR1* mutations, but not healthy volunteers, n=7 healthy volunteers, r= −0.5122, p=0.2398; n=9 *RYR1* mutations, r=0.7511 *p=0.0196.

## Discussion

Herein we describe ER/SR calcium dysregulation promotes exodosis in a human skeletal muscle cell line. Perturbations to ER/SR calcium contribute to the pathophysiology of vast disease states [25], thus presenting a substantial need for the identification of clinical biomarkers and calcium-stabilizing therapeutics.

Despite small patient populations, there are approximately 7,000 rare diseases affecting 350 million patients globally, with a cumulative estimation being greater than HIV and cancer combined [26]. Most rare diseases still lack treatment despite recent efforts to advance rare disease research and accelerate drug discovery [26]. Towards the development of therapeutic modalities, protein-based therapeutics specific to extracellular targets are a viable option [26]. Having the ability to modulate extracellular targets first requires their identification, thus highlighting the translational appeal of investigating exodosis.

Skeletal muscle cells exhibited an exodosis phenotype upon ER/SR calcium dysregulation. While the consequences for redistributing the resident ER/SR proteins from intracellular to extracellular space need to be determined, circulating ERS proteins have been identified in human plasma from a variety of diseases associated with ER calcium dysregulation [27–30]. Many extracellular ERS proteins identified in our analyses are molecular chaperones that interact with, stabilize, or assist other proteins reach functional conformation (e.g., HSP, PDI, CALR) [31]. Protein disulfide isomerases (PDIs), specifically, are a family of ER/SR-resident proteins responsible for the maintenance of disulfide bonds, thus promoting naïve protein conformation [32]. The *PDI* gene family encompasses 21 genes that all contain a thioredoxin domain, but are quite diverse in size, expression, and function [33]. Increased extracellular PH4B, PDIA5, PDIA4, and PDIA3 were identified in mass spectrometry analysis (Supplemental Figure 1B). Extracellular PDIA3 has been implicated as a mediator of myoblast differentiation and fusion during muscle regeneration [34]. Muscle regeneration is integral to the restoration of physiological function and when impaired, as often observed in myopathies, can result in weakened function [34]. Moreover, PDIs are upregulated upon unfolded protein response (UPR) activation, serving as a molecular chaperone set forth to alleviate protein misfolding through rearrangement of incorrect disulfide bonds via isomerase activity or through the degradation of misfolded proteins via ER associated degradation (ERAD) [35, 36]. Relocation of molecular chaperones to the extracellular environment suggests a loss of intracellular proteins critical to maintaining proteostasis, perhaps exacerbating disease progression and symptoms experienced by RYR1-RM- affected individuals.

Liver carboxylesterases (CES1 and CES2) are ERS proteins that undergo exodosis in response to ER calcium depletion [37]. Our lab previously developed an extracellular esterase assay as an indicator of ER calcium depletion in the liver caused by high fat diet [38]. Although predominantly expressed in liver, CES2 has also been detected in bladder, heart, small intestine, and skeletal muscle [39]. The function of CES2 in skeletal muscle has not been studied. Here, using mass spectrometry, we found that CES2 protein is increased in the extracellular space following ER/SR calcium depletion. We also detected increased CES1/2 activity in the media of thapsigargin treated skeletal muscle cells. Our data suggest that CES2 undergoes exodosis in skeletal muscle cells and further studies into the function of CES2 in skeletal muscle are warranted.

One ERS protein in particular, MANF is an ER stress inducible protein that exhibits vast trophic activity among various tissues and pathological conditions [40]. The secretion of MANF is triggered by ER stress associated with ER calcium depletion [8, 41]. We observed an increase in extracellular MANF levels following ER/SR calcium in human skeletal muscle cells. Mutations in *RYR1* can lead to prolonged ER stress and UPR activation [42]. Prolonged ER stress can contribute to muscle atrophy, inflammation, insulin resistance, and disrupted proteostasis [43]. Increased release of pro-inflammatory cytokines IL-6 and IL-1 from cultured myotubes and B-lymphocytes has been observed in MH-causing *RYR1* mutations [44, 45], suggesting a potential interplay with the immune system. There is a fine balance between local, transient inflammation that stimulates pro-myogenic muscle repair and wide-spread, chronic inflammation that diminishes regenerative capability [46]. While the function of MANF in skeletal muscle cells is limited, its anti-inflammatory effects are well documented [40]. Our data from human patients with *RYR1* mutations show a correlation of age and increased plasma levels of MANF. Whether the elevated MANF is indicating ER/SR calcium depletion in skeletal muscle is not clear, however, human patients without the mutations did not see an age-related increase in plasma MANF levels. Also, decreased circulating MANF levels have been reported in human and rodent sera in an age-dependent manner [47], contrary to our observation of age-dependent MANF increase specific to RYR1-RM-affected individuals. While MANF displays biomarker potential for the RYR1 community, we recognize the limited sample size in the current study, emphasizing the need for more rare disease natural history studies. However, our collective data support MANF as a potential biomarker of skeletal muscle diseases associated with ER/SR calcium dysregulation. Additional studies are needed to further examine the relationship of MANF and RYR1-related myopathies.

In addition to identifying ERS-containing proteins, several non-ERS proteins were also detected outside of the cell following ER/SR calcium depletion. For example, SEPN1 was the overall protein (ERS and non-ERS) that showed the highest increase in media of cells treated with thapsigargin. This is largely due to no detectable SEPN1 protein in some vehicle samples and detectable signal in Tg-treated samples. SEPN1 is a type II ER transmembrane protein with reductase activity sensitive to ER calcium fluctuations [48]. SEPN1 has been proposed as an intermediary between calcium sensing and calcium refilling through SERCA2 interactions. Single amino acid mutations in SEPN1 are associated with SEPN1-related myopathy (SEPN1-RM), chronic ER stress, altered calcium affinity, and impaired SEPN1 conformational change [48]. BioGRID analysis of SEPN1 indicated interactions with ERS proteins, TNRC5 and OS9, both of which displayed a 4.247-fold and 1.15-fold increase in response to ER/SR calcium depletion, respectively (Supplemental Figure 1B). Perhaps SEPN1 is cleaved during ER/SR calcium depletion and its interaction with ERS proteins is sufficient to confer its release from the cell, ultimately exacerbating alterations to proteome composition. To our knowledge, this is the first to report the presence of extracellular SEPN1 in response ER/SR calcium depletion. This may present insight into SEPN1 handling in individuals with SEPN1-RM and poses biomarker potential. Interestingly, SEPN1-RM shares some clinical and histopathologic features with RYR1-RM [49]. Mass spectrometry or biomarker analyses of plasma from SEPN1-RM individuals and healthy volunteers may be warranted to corroborate our *in vitro* findings.

A second non-ERS proteins that was elevated following ER/SR calcium depletion is SIL1. SIL1 is a co-factor of ERS protein, HSPA5 (BiP) [50], which displayed a 5.6115-fold increase following ER/SR calcium depletion (Supplemental Figure 1B). Causative mutations in *SIL1* are associated with the rare genetic disorder Marinesco-Sjögren syndrome (MSS) [51]. MSS is characterized by intellectual deficits, congenital cataracts, and myopathy, with some genotype-phenotype data indicating protein instability, protein aggregation, and ER stress [52]. Proteomic profiling of wild-type and *Sil1*^GT^ mice indicated perturbations to muscle physiology evidenced by decreased expression of proteins associated with insulin receptor signaling and glucose metabolism, along with increased expression of proteins associated with ER stress and UPR activation [51]. Elucidating ER/SR proteome redistribution can provide insight into disease pathology, biomarker identification, and drug discovery. Implementing the examination of exodosis for compound screens could prove beneficial to developing a multi-faceted therapeutic approach.

Bromocriptine was recently described for its anti-exodosis effects [10]. Here we show that pre-treatment with bromocriptine elicits anti-exodosis effects in skeletal muscle cells. Bromocriptine has been used clinically to treat acromegaly and Parkinson’s disease [53, 54]. Until recently, bromocriptine was thought to act solely on D2 dopamine receptors (D2R), but its effects on exodosis appear to work in a D2R-independent manner, with ability to signal through GPCRs [10] Although the exact target by which bromocriptine prevents exodosis is currently unknown [10]. Surprisingly, dantrolene exhibited minimal exodosis attenuation unlike previous reports [8, 10]. This may be due to differences in cell lines and varying expression levels of RYR1.

The use of Tg to deplete ER/SR calcium stores is in skeletal muscle cells is well documented [55–59]. Many cellular model systems of RYR1-RM rely on the transfection of mutant *RYR1* cDNA into HEK293 cells, which lack several components responsible for the regulation of RYR1 function [60]. Furthermore, many myopathies fall under the rare disease category, making it challenging to procure primary tissue containing clinically reported mutations. We certainly recognize Tg treatment alone does not fully recapitulate the pathophysiology of myopathies, the observed results do suggest a therapeutic role of identified compounds and potential biomarkers in pre-clinical models of ER/SR calcium dysregulation.

Further studies are needed to assess the efficacy of these compounds and utility of the biomarkers in the genetic context of RYR1-related myopathies.

## Materials and Methods

### Human Samples

Human samples were obtained from the National Institute of Nursing Research, clinical trial identifier NCT02362425. The protocol was approved by the NIH Combined Neuroscience Institutional Review Board. All participants provided written informed consent.

### Cell Culture

T0034 immortalized human skeletal muscle cell line was purchased from ABM. Cell were maintained in Prigrow III medium (ABM), 10% FBS (Sigma), 1% Pen/Strep (Life Technologies) and grown at 37°C with 5.5% CO2 in a humidified incubator. Applied extracellular matrix (ABM) was used to coat culture vessels for cell line maintenance and experiments.

For mass spectrometry experiments, cells were seeded in ECM coated 15cm plates at 8×10^6^cells/plate. Approximately 72 hours post-plating, media was removed, cells were rinsed with 1X PBS, and replaced with mass spectrometry assay media (150mM NaCl, 5mM KCl, 1mM MgCl_2_, 20mM HEPES, 1mM CaCl_2_, H2O). Cells were treated with vehicle (.5% final DMSO), 20μM bromocriptine (Tocris), or 50μM dantrolene (Caymen Chemical) for 30 mins, followed by vehicle (0.1% DMSO) or 100nM Tg (Sigma). After 8 hours, 20mL of media was collected, centrifuged at 1000rpm for 5 minutes at room temperature to pellet debris. 15mL of supernatant was concentrated using 10,000MW cutoff concentrator conical tubes (Millipore) centrifuged at 4000xg for 40 minutes at 4°C as previously described [8]. Protein concentration was determined by DC assay (BioRad). Mass spectrometry analyses were performed by the Johns Hopkins University Mass Spectrometry and Proteomics Core Facility. Briefly, 50μg of protein was analyzed by TMT pro-16-plex labeling (Thermo Fisher), peptides fractionated by basic reverse phase (bRP) chromatography and analyzed by liquid chromatography tandem mass spectrometry (LC-MS/MS).

### Western blot analysis

Ten micrograms of protein from concentrated media were collected as described above was set aside for western blot analysis. Proteins were separated on 4-12% Bis-Tris NuPage gels (Thermo Fisher) using MOPS running buffer for 50 minutes at 200V. Proteins were transferred to 0.2μM PVDF membrane using iBlot2 (Thermo Fisher), immunoblotted for MANF (Yenzym #2156; 1:500), P4HB (Abcam #2792; 1:500) and scanned using Licor system. For protein stain, gels were stained using Simply Blue Safe Stain (Thermo Fisher) and imaged on Azure gel doc system.

### Immunoprecipitation Assay

Media collected from T0034 cells were immunoprecipitated with magnetic Protein A beads (Sure Beads, BioRad) as previously described [8]. Beads were washed with 1X PBS plus 0.1% Tween 20 (PBS-T) and incubated with 2μg mouse anti P4HB (Abcam #2792). 400μL of media was incubated with beads for 1 hour followed by a series of PBS-T washes as recommended by manufacturer protocol. Samples were eluted in 40μL of 1X LDS (Thermo Fisher), incubated at 70°C for 10 minutes, and 18μL was loaded for western blot as described above.

### Esterase Activity

Methods to measure CES1/CES2 activity have been previously described [37]. Briefly, 20μL of cell culture media was transferred to black walled, clear bottom plate. Equal volume of 100μM of fluorescein di-(1-methylcyclopropanecarboxymethyl ether) substrate diluted in mass spectrometry media at pH 5.0 was added to each well. Fluorescence was measured every minute for 60 minutes at 45°C using BioTek Synergy H2 plate reader (excitation 485nm, emission 528nm).

### MANF Homogenous Time Resolved Fluorescence Assay

MANF HTRF assay (CisBio) was used to determine circulating MANF concentration from 16μL of human plasma samples or cell culture media according to manufacturer’s instructions. Cell culture samples were collected in culture media or mass spectrometry assay media and diluted 1:5 in assay diluent. Samples were read on BioTek Synergy H1 plate reader. 2μL of anti-Human MANF d2 and 2μL of anti-Human MANF Eu Cryptate antibody working solutions were added to wells and incubated together in a sealed, white-walled plate for 4 hours at room temperature. Fluorescence was measured on BioTek Synergy H1 plate reader at 665 nm and 620 nm. Sample readouts were compared to a MANF standard curve to determine MANF concentration in each sample.

### Statistical Analysis

Results were analyzed using Graph Pad Prism 9. Details about tests can be found in figure legends. Proteome Discoverer 2.4 was used for the generation of biological function pie chart. Data are presented as mean ± SEM and analyzed using two-tailed t-tests, 1-way ANOVA with Dunnett’s multiple comparison tests, 2-way ANOVA with Sidak’s multiple comparison test, and Pearson correlation test. Investigator was not blinded to healthy versus RYR1-RM plasma samples but was blinded to personal identifiable information.

## List of Abbreviations

CCD: Central core disease
Ces1/Ces2: Liver carboxylesterase 1 or 2
CFTD: Congenital fiber-type disproportion
CNM: Centronuclear myopathy
D2R: Dopamine 2 receptor
ER/SR: Endoplasmic reticulum/sarcoplasmic reticulum
ERAD: ER associated degradation
ERS: ER retention/retrieval sequence
FDA: Food and Drug Administration
GPCR: G-protein coupled receptor
MANF: Mesencephalic astrocyte derived neurotrophic factor
MH: Malignant hyperthermia
MmD: Multi minicore disease
PDI: Protein disulfide isomerase
RYR1: Ryanodine receptor-1
RYR1-RM: RYR1-related myopathies
SEPN1: Selenoprotein 1
SEPN1-RM: SEPN1-related myopathies
SIL1: Nucleotide exchange factor 1
UPR: Unfolded protein response

## Acknowledgements

We thank the Johns Hopkins University Mass Spectrometry and Proteomics Core for data acquisition.

## Authors’ Contributions

E.S.W, L.K.G., B.K.H conceived and executed experiments. E.S.W and L.K.G. analyzed data, E.S.W, L.K.G and B.K.H. wrote manuscript. K.G.M. provided clinical expertise, manuscript review, and human plasma samples.

---

**Supplemental Figure 1.**
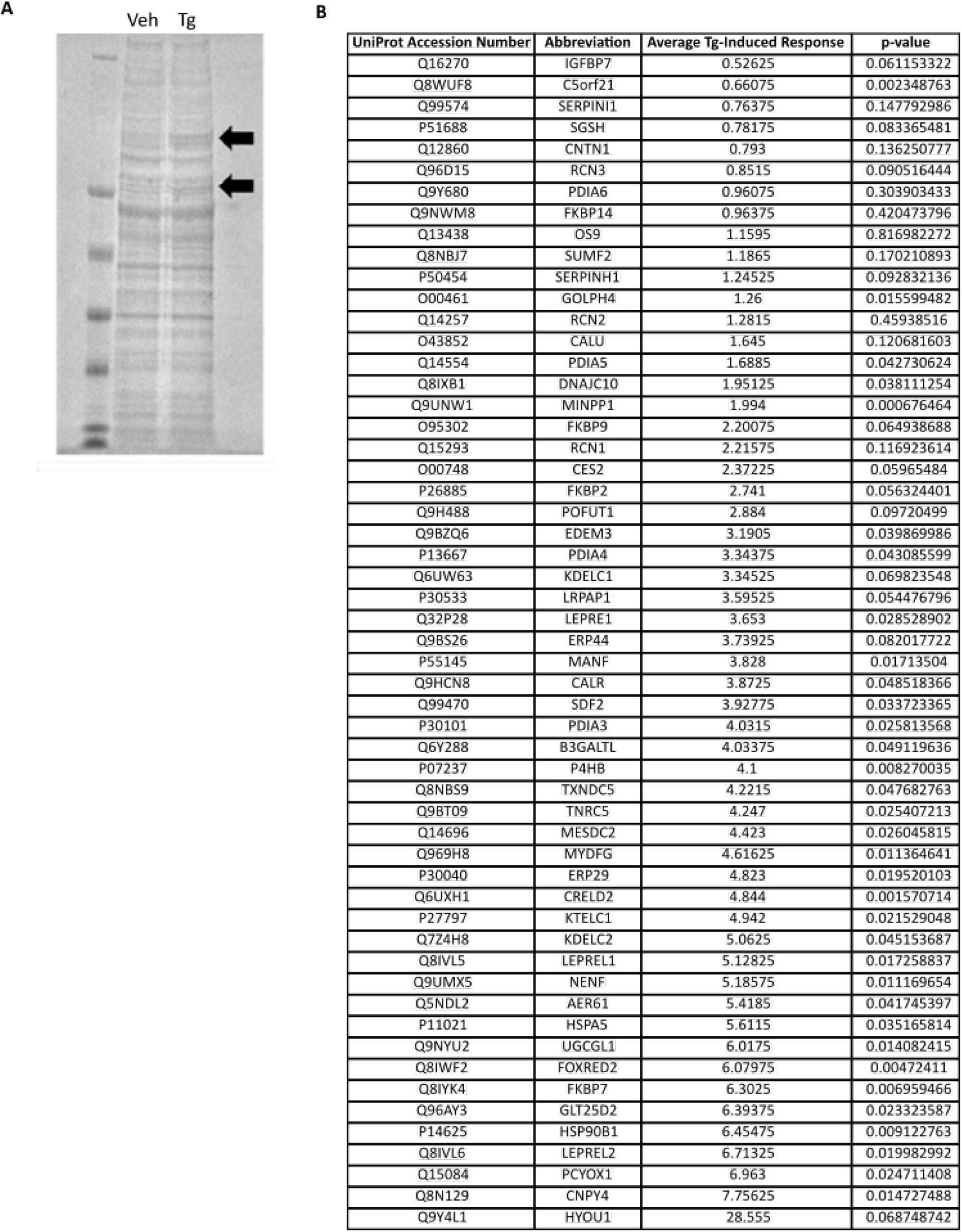
Identification of extracellular ERS proteins. A) Blue stain gel of concentrated media from skeletal muscle cells treated with vehicle or 100nM Tg for 8 hours. Arrows indicate qualitative protein increases. B) Table of extracellular ERS proteins identified by mass spectrometry, UniProt accession number, and average Tg-induced response, p-values listed, 2-tailed t-test, vehicle normalized abundance vs Tg normalized abundance.

**Supplemental Figure 2.**
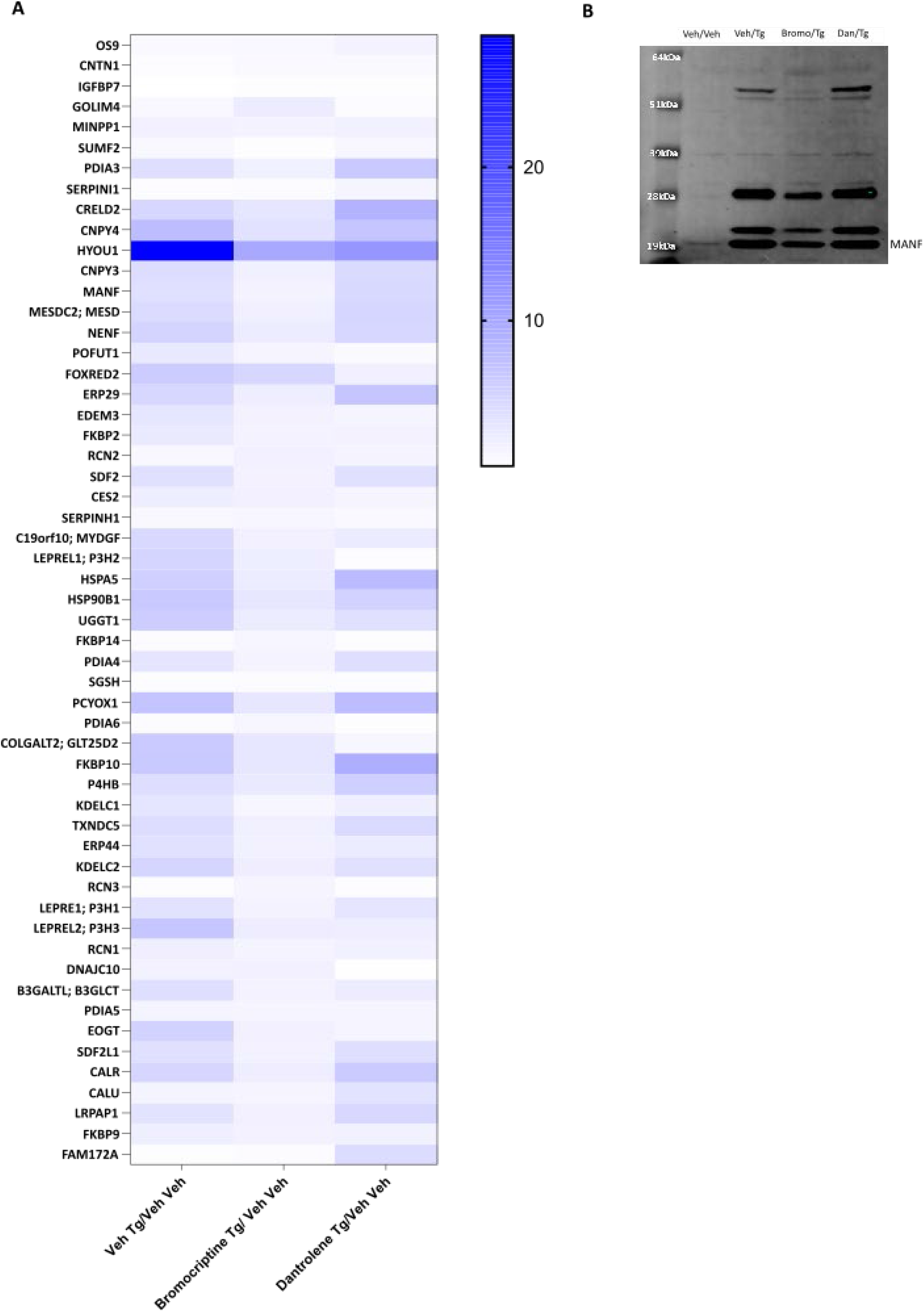
Extracellular ERS proteins pre-treated with bromocriptine or dantrolene. A) Heat map depicting changes in extracellular ERS protein content when pre-treated with 20μM bromocriptine, or 50μM dantrolene 30 minutes prior to 100nM Tg for 8 hours. B) Western blot of concentrated media collected from T0034 skeletal muscle cell line pre-treated with vehicle, 20μM bromocriptine, or 50μM dantrolene, mean ± SEM, n=2/treatment groups.

